# Two functionally distinct Purkinje cell populations implement an internal model within a single olivo-cerebellar loop

**DOI:** 10.1101/2021.05.09.443096

**Authors:** Dora E. Angelaki, Jean Laurens

**Affiliations:** Center for Neural Science and Tandon School of Engineering, New York University, NY, USA; Ernst Strüngmann Institute (ESI) for Neuroscience in Cooperation with Max Planck Society, Frankfurt, Germany

## Abstract

Olivo-cerebellar loops, where anatomical patches of the cerebellar cortex and inferior olive project one onto the other, form an anatomical unit of cerebellar computation. Here, we investigated how successive computational steps map onto olivo-cerebellar loops. Lobules IX-X of the cerebellar vermis, i.e. the nodulus and uvula, implement an internal model of the inner ear’s graviceptor, the otolith organs. We have previously identified two populations of Purkinje cells that participate in this computation: Tilt-selective cells transform egocentric rotation signals into allocentric tilt velocity signals, to track head motion relative to gravity, and translation-selective cells encode otolith prediction error. Here we show that, despite very distinct simple spike response properties, both types of Purkinje cells emit complex spikes that are proportional to sensory prediction error. This indicates that both cell populations comprise a single olivo-cerebellar loop, in which only translation-selective cells project to the inferior olive. We propose a neural network model where sensory prediction errors computed by translation-selective cells are used as a teaching signal for both populations, and demonstrate that this network can learn to implement an internal model of the otoliths.

## Introduction

Theories developed over the last decades (Ito, 2006; Kawato, 1999; Wolpert et al., 1998a, 1998b) have proposed that the cerebellum implements forward internal models that predict sensory inflow based on internal representations of the world and our body. Sensory predictions are then compared to actual sensory afference. In the event of mismatches, the resulting sensory prediction errors drive corrective feedback mechanisms to update internal representations and guide perception and action. On a longer time scale, these errors drive learning mechanisms to acquire or calibrate the internal models (Herzfeld et al., 2018; Kimpo et al., 2014; Lisberger, 1988; Nguyen-Vu et al., 2013).

Cerebellar computations are implemented by olivo-cerebellar loops (**Fig. 1A**) (Apps et al., 2018; Apps & Garwicz, 2005; Chaumont et al., 2013; De Zeeuw et al., 2011; Ozden et al., 2009; Sugihara & Quy, 2007), within which a group of Purkinje Cells (PCs) in the cerebellar cortex project simple spikes (SS) to a group of cells in the cerebellum’s output nuclei (the deep cerebellar nuclei, DCN, and vestibular nuclei, VN). These nuclei project throughout the nervous system to control behaviour, and to a group of Inferior Olive (IO) neurons that projects back to the cerebellar cortex. IO neuron influence on PCs induces Complex Spikes (CS) that act as teaching signals to drive cerebellar learning (Herzfeld et al., 2018; Kimpo et al., 2014; Lisberger, 1988; Nguyen-Vu et al., 2013). PCs within an olivo-cerebellar loop are anatomically clustered in sagittally oriented microzones of several hundred microns in length and tenths of microns in width (Kostadinov et al., 2019; Ozden et al., 2009; Valera et al., 2016). An olivo-cerebellar loop can be formed by multiple microzones that receive similar projections from the IO, and collectively form a multizonal microcomplex (Apps & Garwicz, 2005; Cerminara et al., 2020): in this study we will use the term ‘microcomplex’ to refer to a set of PCs receiving identical IO projections, and ‘loop’ to refer to the network of cortical, nuclear and IO neurons communicating with a microcomplex.

**Figure 1:**
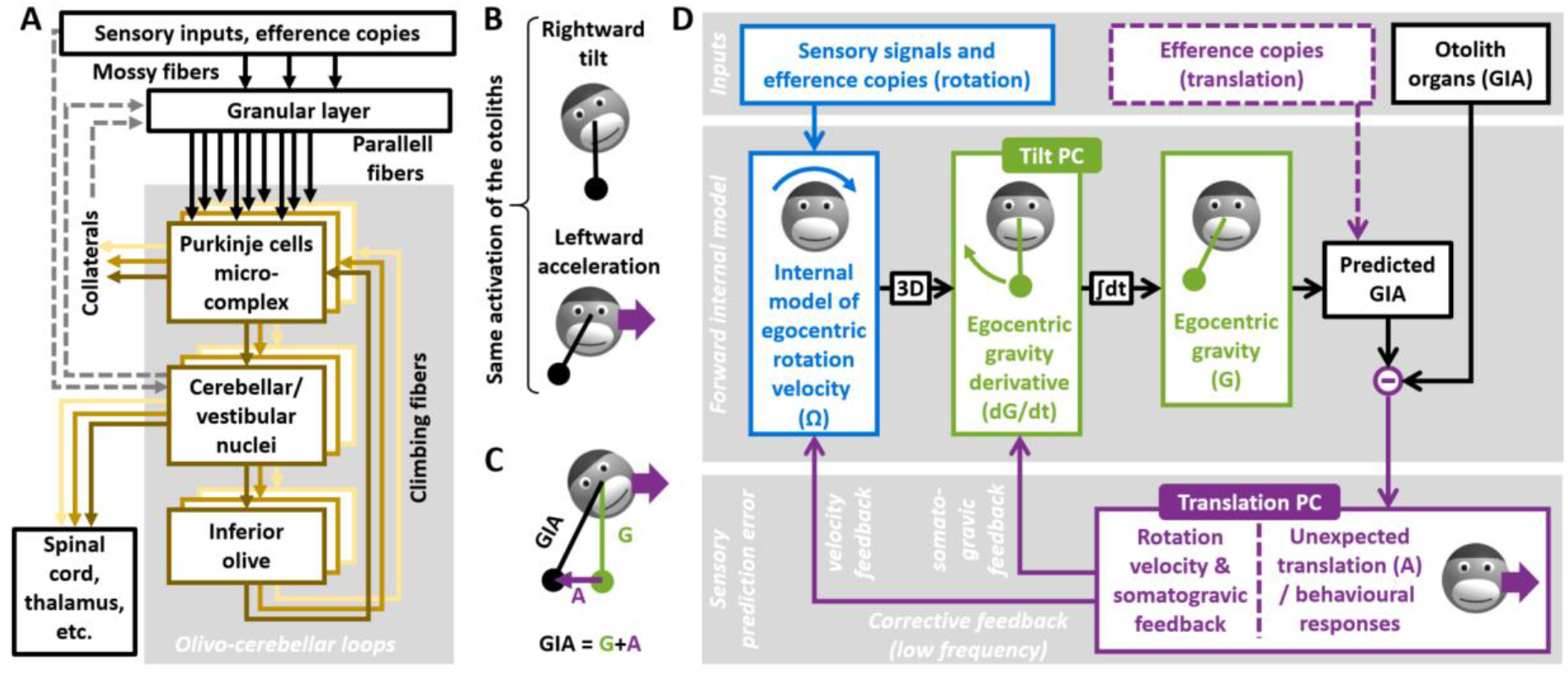
Olivo-cerebellar loops, and internal model computations for processing otolith signals. **A:** Neural pathways and information processing in olivo-cerebellar loops. Sensory inputs and efference copies reach the cerebellum though mossy fibers and are processed in the granular layer. Granule cells convey this information to PCs though parallell fibers. PCs are anatomically clustered in microzones, and several microzones participating to a single olivo-cerebellar loop form a microcomplex. Further pathways exist in the vestibular circuitry: primary afferents also reach the vestibular nuclei, which project to the granular layer, and PCs may project collaterals to the granular layer. **B:** Ambiguity of the otolith organs. The otolith organs are analogous to a pendulum, whose position is sensed in egocentric head coordinates. Rightward head tilt or leftward acceleration cause a rightward deviation of the pendulum relative to the head, resulting in an identical activation of the otoliths. **C:** Mathematical formulation of the ambiguity. The otoliths sense the gravito-inertial acceleration (GIA), expressed as GIA=G+A where G is the gravity vector and A a vector opposite to the linear acceleration (this convention is chosen for clarity purposes). The brain may resolve the ambiguity by tracking G, and computing A by subtraction (A=GIA-G). **D:** Internal model computations for otolith information processing. See text for description.

Studies to date have pioneered the ‘microcomplex’ as a fundamental unit of cerebellar computation, e.g. during saccadic eye movements (Herzfeld et al., 2015, 2018), tactile reflexes (Apps & Garwicz, 2005; Cerminara et al., 2020; Ekerot et al., 1991; Garwicz et al., 1998) or cognitive tasks (Kostadinov et al., 2019). This has led to the notion that identifying PCs that receive identical IO inputs (i.e. participate in the same microcomplex) allows parsing the cerebellar cortex into elementary computation units (Herzfeld et al., 2015, 2018; Shadmehr, 2020). However, how to map such multivariable computations onto olivo-cerebellar loops raises fundamental questions. One possibility is that each variable is represented by a different microcomplex such that multivariable computations are implemented by parallel loops, each computing one variable. Alternatively, it is also possible that PCs encoding fundamentally distinct variables may exist in a single microcomplex. Such a finding would depart from the traditional view where one loop computes one variable and suggest that individual microcomplexes can perform sequences of operations: Functionally distinct PCs perform distinct computations using common teaching signals.

To distinguish between these two hypotheses we take advantage of a multivariable cerebellar computation based on an internal model of self-motion, already widely studied in the literature (**Fig. 1B-D**) (Borah et al., 1988; Bos & Bles, 2002; Glasauer & Merfeld, 1997; Karmali & Merfeld, 2012; Laurens & Angelaki, 2011, 2017; Laurens & Droulez, 2007; Merfeld, 1995; Oman, 1982; Ormsby & Young, 1977; Zupan et al., 2002). A unique advantage of this system is the ability to map complex, but well-understood, algorithmic computations implementing an internal model of the inner ear’s inertial motion sensors, the otolith organs, into a cerebellar circuit that includes lobules X and IX of the cerebellar vermis (Nodulus and Uvula; NU) (Laurens et al., 2013a, 2013b; Laurens & Angelaki, 2020; Stay et al., 2019; Yakusheva et al., 2007, 2008, 2013).

Specifically, the otolith organs sense the sum of gravitational (G) and linear accelerations (A), which are physically indistinguishable (Einstein, 1907), in head coordinates (**Fig. 1 B,C**). The otolithic signal is therefore inherently ambiguous. Nevertheless, this ambiguity can be resolved by using additional sensory information and motor inference copies to predict the two components of otolith activation, gravity (G) and translational acceleration (A). On the one hand, the gravitational component G can be predicted by tracking head rotation relative to gravity (**Fig. 1D**, green). A portion of the head’s internal model of motion (**Fig. 1D**, blue; not developed here for simplicity; (see (Karmali & Merfeld, 2012; Laurens & Angelaki, 2011, 2017) for details), senses head rotation velocity (Ω) in an egocentric frame of reference. The internal model converts Ω into allocentric velocity relative to gravity (block marked ‘3D’ in **Fig. 1D**), which is equivalent to the derivative of gravity in head coordinates (dG/dt). This signal is integrated over time (block marked ‘∫’ in **Fig. 1D**) to estimate the gravity vector in head coordinates (G). On the other hand, head translation may be derived directly from motor efference copies during active translation (**Fig. 1D**, violet, broken lines), but is unpredictable during passive movements.

Altogether, the internal model can predict otolith signals during active tilt and translations (based on motor commands), or during passive tilt (based on rotation signals). Thus, otolith prediction errors occur during passive translations, or if tilt signals are erroneous, which can occur because of sensory noise or incorrect rotation signals from the canals. Since these tilt errors are generally smaller and scarcer, the brain preferentially interprets otolith prediction errors as translation. Accordingly, otolith prediction errors induce a perception of translation and the corresponding stabilizing eye movements, irrespective of whether the prediction error originates from an actual translation or an artificially generated incorrect canal signal (Angelaki et al., 1999; Hess & Angelaki, 1999; Khosravi-Hashemi et al., 2019; Merfeld et al., 1999). Otolith prediction errors also trigger low-frequency feedback (**Fig 1D**, violet) that gradually correct the underlying rotation signals and tilt estimates.

Based on SS responses exclusively, two populations of PCs were identified that perform distinct steps in the internal model’s computation. First, translation-selective cells (**Fig. 1D**s) encode otolith predictions error (Laurens et al., 2013a, 2013b; Laurens & Angelaki, 2020; Stay et al., 2019; Yakusheva et al., 2007, 2008, 2013). These cells respond selectively to passive translation, indicating that they (i) receive otolithic inputs, (ii) are cancelled by tilt signals originating from rotation sensing (**Fig. 1D**; (Laurens et al., 2013b) and (iii) encode sensory prediction errors that result from artificial canal stimulation (Laurens et al., 2013a). Critically, the responses of translation-selective cells in the VN and DCN are attenuated during active head translations (Carriot et al., 2013; Mackrous et al., 2019), a finding that confirms that the internal model uses efference copies to predict otolith signals.

Second, another PC type in the NU encodes tilt velocity (**Fig. 1D**, tilt-selective cells; (Hernández et al., 2020; Laurens et al., 2013b; Laurens & Angelaki, 2020; Stay et al., 2019). These cells modulate more during tilt than translation in phase to tilt velocity (Laurens & Angelaki, 2020). Importantly, 3D motion stimuli have revealed that these cells encode transformed rotation signals (dG/dt), and not egocentric rotation velocity (Ω) (Laurens et al., 2013b).

Despite a good understanding on the properties of SS responses, little is currently known about CSs, which are fundamental for understanding the organisation of the corresponding cerebellar circuits. Previous CS studies were limited to rotation stimuli (Barmack & Shojaku, 1995; Fushiki & Barmack, 1997; Kitama et al., 2014; Yakhnitsa & Barmack, 2006), or only characterized translation-selective cells (Yakusheva et al., 2010). A crucial, yet unanswered, question is whether CS firing is different in tilt-selective and translation-selective cells: this would imply that there are two distinct cerebellar loops. Alternatively, if tilt-selective and translation-selective cells exhibit similar CS firing, they may comprise a single loop using the same teaching signals.

Here, we analysed the CS firing of both tilt- and translation-selective cells during combinations of tilt and translation stimuli, as well as 3D motion (Laurens et al., 2013a, 2013b). Surprisingly, we found that the CS firing of both cell types is identical, and occurred specifically during translation. This indicates that the teaching signal to both cell types is driven by otolith prediction error, which is the output of the internal model implemented by the NU. We interpret these findings in the context of a previously proposed learning rule (Dean et al., 2002, 2010; Dean & Porrill, 2014), and validate our interpretation by simulating a neural network model that learns to discriminate tilt from translation.

## Results

We analysed CSs of 66 out of 211 Purkinje cells recorded in lobules IX-X of the cerebellar vermis (Laurens et al., 2013a, 2013b) that could be identified consistently across trials and followed by a pause in SS activity for at least 10 ms. Neurons were recorded during sinusoidal translation (**Fig. 2A**, left) and tilt (**Fig. 2A**, middle) at 0.5 Hz (Angelaki et al., 1999, 2004; Laurens et al., 2013a, 2013b; Shaikh et al., 2005; Yakusheva et al., 2007, 2008, 2013), which activated the otoliths identically (**Fig. 2A**, GIA). A few cells were also recorded during: (1) out-of-phase tilt and translation (tilt-translation, **Fig. 2A**, right), where linear acceleration and tilt cancel each other, such that the otoliths are not activated but the canals sense velocity; and (2) in-phase tilt + translation (not represented in figures; see (Laurens et al., 2013b; Laurens & Angelaki, 2016)). We used a spatio-temporal tuning model (Laurens et al., 2013b; Laurens & Angelaki, 2016) together with a bootstrap test to classify cells as translation-selective (larger response to translation), tilt-selective (larger response to tilt), GIA-selective (same response to tilt and translation, similar to otolith afferents), composite (cells who could not be classified in one of these categories) or non-responsive.

**Figure 2:**
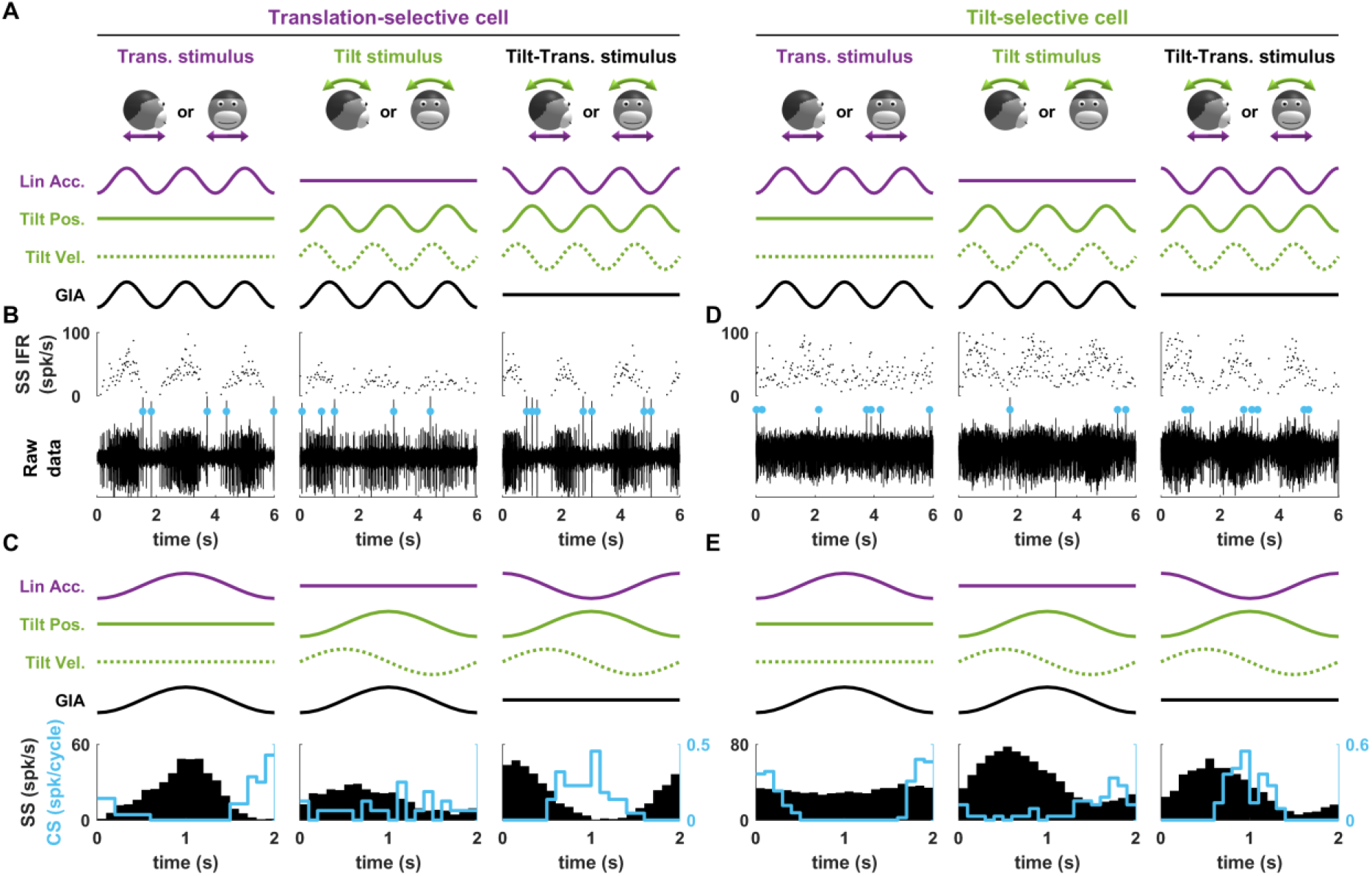
Representative Purkinje cells during tilt/translation. **A:** illustration of the motion stimuli. Violet and solid green curves: inertial acceleration and tilt position; the sum gives the GIA (black). Tilt velocity is indicated by broken green curves. **B:** Spiking activity of a translation-selective cell. Bottom traces show the raw extracellular voltage. CSs are marked by cyan dots. Upper traces show instantaneous firing rate (IFR) of the SSs. **C:** Average firing histograms of SSs (black) and CSs (cyan). **D,E:** Spiking activity and average firing of a representative tilt-selective cell (layout as in B,C).

### Example cells

Responses of example tilt- and translation-selective neurons are shown in **Fig. 2**. The translation-selective cell (**Fig. 2B,C**) shows vigorous SS response during translation (**Fig. 2B,C**, left), but not during tilt (**Fig. 2B,C**, middle). During translation, the cell fired CSs during the trough of the SS response (**Fig. 2B,C**, left, marked by cyan dots). In contrast, the phase-locked firing was weaker during tilt (**Fig. 2B,C**, middle). During tilt-translation (**Fig. 2B,C**, right), SSs and CSs maintained their phase relationship relative to the translational component of the stimulus, as expected if they were both driven by translation.

The second example neuron (**Fig. 2D,E**) is representative of tilt-selective cells (Laurens et al., 2013b; Laurens & Angelaki, 2020; Stay et al., 2019): SS modulation was higher during tilt compared to translation (**Fig. 2D,E**, middle versus left). Consistently, groups of 2-3 CSs occurred at regular phases during each cycle of translation (**Fig. 2D**, left), such that a clear CS modulation occurred during translation (**Fig. 2E**, left). In contrast, CS modulation was weaker during tilt (**Fig. 2E**, middle). During tilt-translation (**Fig. 2D,E**, right), when SSs occurred during tilt (**Fig. 2E**, right versus middle), CS modulation maintained its phase with respect to the translational component of the stimulus. Thus, this cell’s CS firing (**Fig. 2E**) was locked to head translation, and conspicuously similar to that of the translation-selective cell (**Fig. 2C**).

### SS and CS response gains

Consistent with previous studies (Laurens et al., 2013b, 2013b; Laurens & Angelaki, 2020; Stay et al., 2019), cells were classified based on their SS response gain to tilt and translation, computed along the preferred direction (PD) and expressed in identical units of spk/s/G. By definition, the gains of 18/62 cells (29%) translation-selective cells appear below the diagonal (**Fig. 3A**, violet) since they respond more to translation, spanning a range of 100 to 1000 spk/s/G. The gains of 24/62 (39%) tilt-selective cells appear above the diagonal (**Fig. 3A**, green), spanning 20 to 300 spk/s/G during tilt, but orders of magnitude smaller during translation. GIA-selective and composite cells (20/62, 32%) lie close to the diagonal. These proportions and ranges of response gains resemble those reported by the broader cell population (Laurens et al., 2013b), indicating that the cells analysed here are representative of the full population. Furthermore, they are similar to the population responses of subsequent studies using stimuli based on Gaussian (rather than sinusoidal) temporal profiles (Laurens & Angelaki, 2020) or recorded in mice (Stay et al., 2019).

**Figure 3:**
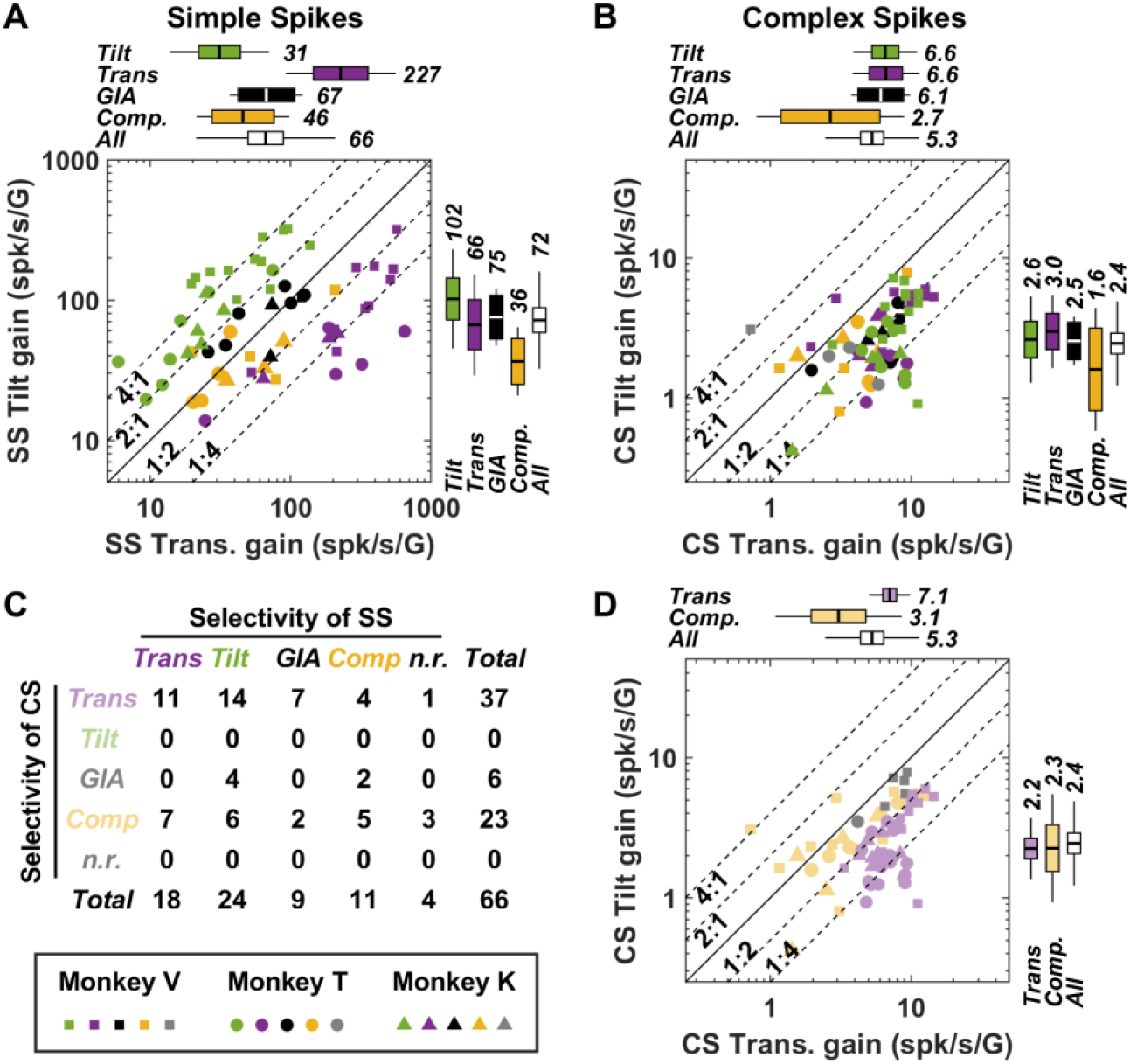
Response modulation and classification of the SS and CS firing across the population of PCs. **A:** Tilt versus translation response gain of SS firing. Cells are color-coded based on their classification (green, violet, back and yellow: tilt-, translation-, GIA-selective and composite). Marker shapes indicate the animal in which cells were recorded (see legend on lower left corner). The boxes and whisker plots represent geometrical average (box center), 95% confidence interval (box) and standard deviation (whiskers) of the gain for each cell type and all cells together (white). Broken black lines parallel to the diagonal represent the level at which tilt response gain is 4x, 2x, 1/2x and 1/4x the translation response gain. **B:** Tilt versus translation response gain of CS firing. Cells are color-coded based on their SS classification, i.e. as in A. **C:** Contingency matrix between the classification of SS and CS response sensitivity. **D:** Tilt versus translation response gain of CS firing, as in B, but with cells classified based on their CS response (green, violet, grey and yellow: tilt-, translation-, GIA-selective and composite).

We next examined the modulation gain of CS. In **Fig. 3B**, neurons are color-coded based on the selectivity of their SS response, i.e. as in **Fig. 3A**. Remarkably, most neurons, including all tilt-selective cells (green), appeared below the diagonal. Thus, like the examples in **Fig. 2**, the CS modulation of both translation- and tilt-selective cells was higher during translation than tilt. In fact, the average CS modulation of tilt-, translation- and GIA-selective cells were identical during translation (6.6, 6.6 and 6.1 spk/s/G respectively, **Fig. 3B**, upper box plots; p=0.79, Krusal-Wallis non-parametric ANOVA) and also during tilt (2.6, 3 and 2.5 spk/s/G respectively, **Fig. 3B**, rightward box plots; p=0.65, Krusal-Wallis non-parametric ANOVA). CS response gains appeared weaker and more variable in composite cells (**Fig. 3B**, yellow boxes) and in cells whose SS didn’t exhibit a significant modulation (**Fig. 3B**, grey markers).

To evaluate whether CS modulation is significant on a cell-by-cell basis, we used the same classification method used for SSs. We found that the majority of CS responses (37/66, 56%, **Fig. 3C**) was independently classified as translation-selective, including most (14/24, 58%) cells that were classified as tilt-selective based on their SSs. These cells appear in violet in **Fig. 3D**. CS modulation was similar during tilt and translation in a few cells (6/66, 9%, **Fig. 3C**; grey markers in **Fig. 3D**). In the rest of the population of cells (23/66, 35%), CS were classified as composite, indicating that they responded to combinations of tilt and translation (**Fig. 3C**; yellow markers in **Fig. 3D**). Translation responses were still larger than tilt responses in the majority (18/23) of these cells. Remarkably, no CS response was classified as tilt-selective. This analysis confirms that CS are generally modulated during translation and not during tilt, regardless of the selectivity of SS responses.

### Spatio-temporal relationships of SS and CS firing

We next investigated whether SS and CS responses are spatially and temporally matched. All cells were recorded along the forward-backward and lateral directions, allowing us to reconstruct the neuron’s tuning curve along all directions (see (Green et al., 2005; Laurens & Angelaki, 2016) for details) and determine the direction along which its response is maximal (PD). These PD are determined separately for SSs and CSs; and we test whether they are aligned in **Fig. 4A**. Note that SSs and CSs may occur along the same axis, but in anti-phase (e.g. as in **Fig. 2C**). In this case, it is equivalent to state that they have similar PD and opposite phase, or that they have similar phase and opposite PD. We adopt the former convention here: as a consequence, the difference in PD between SSs and CSs is never higher than 90°, and the corresponding area is blacked out in **Fig. 4A**. Note also that, since CSs are modulated during translation in tilt-selective cells, we compare the PD and phase of SSs during tilt to the PD and phase of CSs during translation in tilt-selective cells. For all other cell types, we compare SS and CS responses during translation. Note also that PDs are computed relative to the direction of the GIA, which is the stimulus activating the otoliths. For instance, a rightward tilt and leftward acceleration activate the otoliths in the same manner (**Fig. 1B**) and therefore correspond to the same PD.

**Figure 4:**
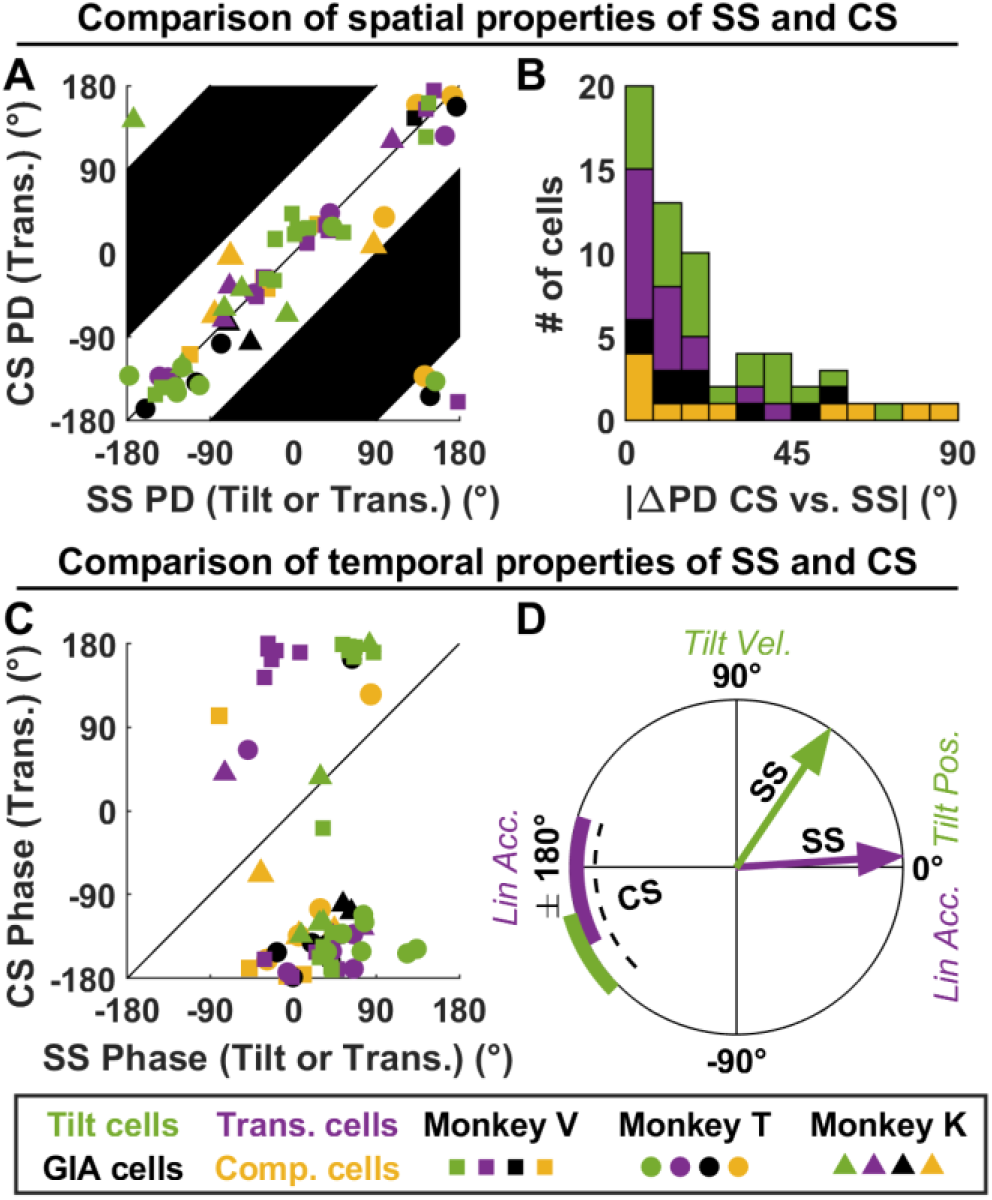
Spatio-temporal comparison of SS and CS responses. **A:** Comparison of the PD of SSs (during tilt in tilt-selective cells and translation in other response groups) and CSs (during translation in all groups). Note that, by convention, the PD of SS and CS are never more than 90° apart (see text; the corresponding areas are marked in black). **B:** Histogram of the PD differences, measured as in A. **C:** Comparison of the response phase of SSs (during tilt in tilt-selective cells and translation in other response groups) and CSs (during translation in all groups) during motion along the PD (defined as in A). **D:** Average response phase of SS (arrow) and CS (sectors representing confidence intervals) in tilt- and translation-selective cells.

We found that the spatial properties of SS and CS were closely aligned. Indeed, their PDs clustered tightly along the diagonal in **Fig. 4A**. To measure how closely the PDs of SSs and CSs align, we computed the absolute difference between them (**Fig. 4B**): this difference can range between 0° (when PDs are aligned) and 90° (when they are orthogonal), and would be distributed uniformly if the PDs of SSs and CSs were independent. We found that this difference was concentrated close to 0° (**Fig. 4B**; median: 14°, [10 19] CI; Kolmogorov-Smirnov test against uniform distribution: p<10^-10^), which confirm that the PDs of SSs and CSs typically align closely.

We next examined the response phase of SSs and CSs. In line with our findings in (Laurens et al., 2013b; Laurens & Angelaki, 2020), the SS response phase of translation-selective cells was close to peak acceleration (**Fig. 4C**, violet; **Fig. 4D**, violet arrow), and that of tilt-selective cells was close to tilt velocity (**Fig. 4C**, green; **Fig. 4D**, green arrow). In contrast, we found that the CS response phase during translation clustered tightly close to −180° in both translation-selective cells (**Fig. 4C,D**; mean: −175°, [−198 −152] CI) and tilt-selective cells (**Fig. 4C,D**; mean: −154°, [−173 −135] CI). This confirms that the CS response of the entire population is homogenous in term of response phase, and identical in tilt- and translation-selective cells.

### The tilt/translation discrimination microcomplex

Previous studies (Herzfeld et al., 2015; Shadmehr, 2020) have proposed that groups of PCs within a microcomplex, i.e. group of PCs that receive similar IO inputs, form a unit of cerebellar computation. Our results indicate that microcomplexes in the NU are formed by mixtures of tilt-, translation-, GIA-selective and composite PCs. In the next analysis, we pooled our data to compute CS and SS responses of average PCs within a NU microcomplex.

To do so, we computed the PD of each cell’s CS, and computed the SS and CS firing histograms across all trials collected within ±45° of the PD. This allowed us to ‘spatially align’ the firing of PCs with various PDs and to average their CS and SS responses. In agreement with **Fig. 3, 4**, we found that the CS response profile of translation-, tilt- and GIA-selective cells are highly similar (**Fig. 5B-D**, cyan). Furthermore, the combined SS activity of translation-selective PCs followed the response pattern of a translation-selective PC (**Fig. 5B**, black; compare e.g. with **Fig. 2C**). This indicates that translation-selective PCs that share similar IO inputs (and thus participate to a common microcomplex), also have similar SS firing, such that the population as a whole can be seen as a single translation-selective PC, a concept that has been described as ‘super-PC’ (Apps et al., 2018). Likewise, the population response of tilt-selective PCs (**Fig. 5C**) is representative of a single ‘super’ tilt-selective PC. In contrast, the SS response modulation of GIA-selective and composite cells was modest, indicating that these groups may not form coherent populations.

**Figure 5:**
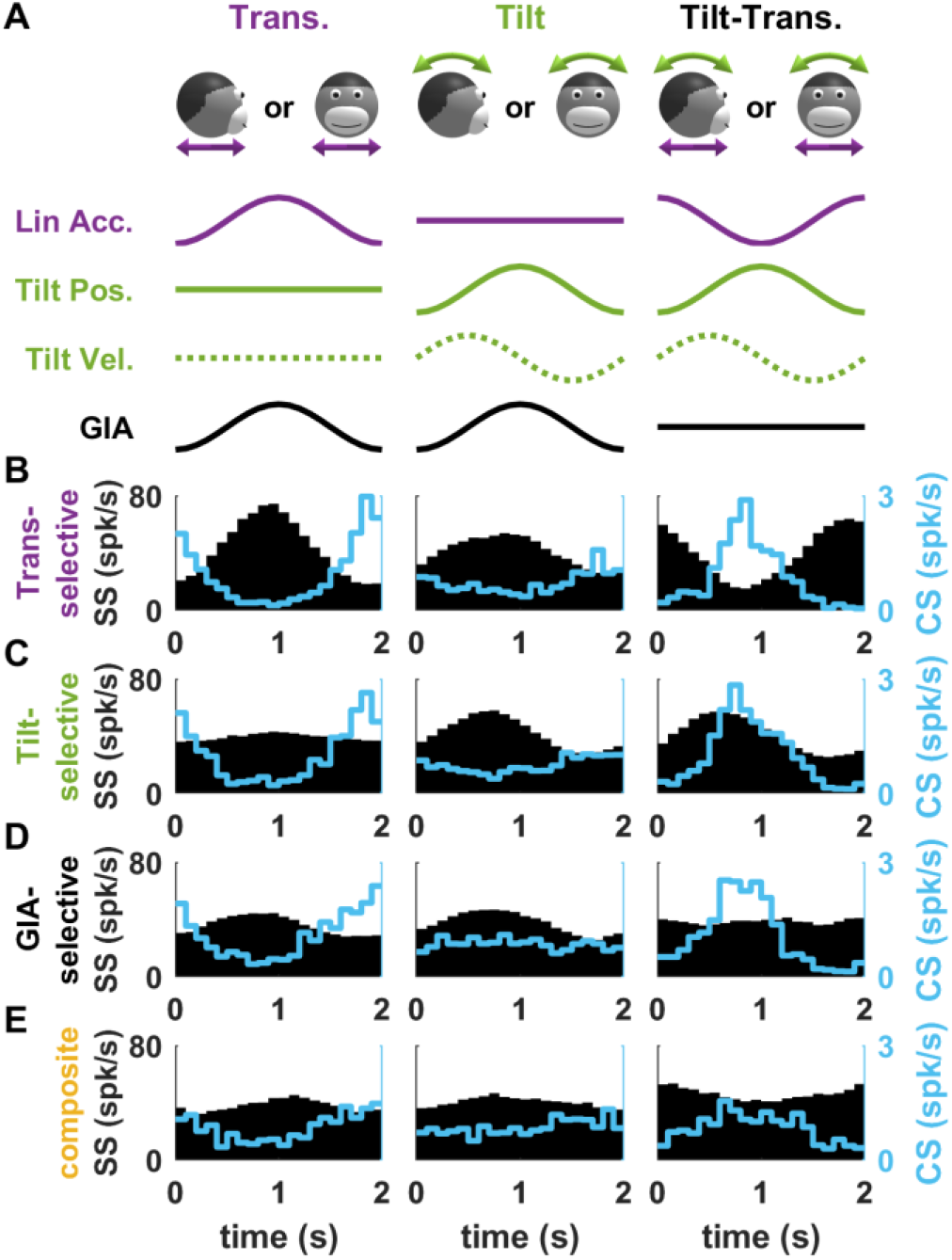
Average SS and CS firing histograms of PCs belonging to a common microcomplex. **A:** Illustration of the motion stimuli and variables. **B-E:** Firing histograms of PCs belonging to all response groups. Black histograms: SS firing. Blue histograms: CS firing.

**Figure 5S1:**
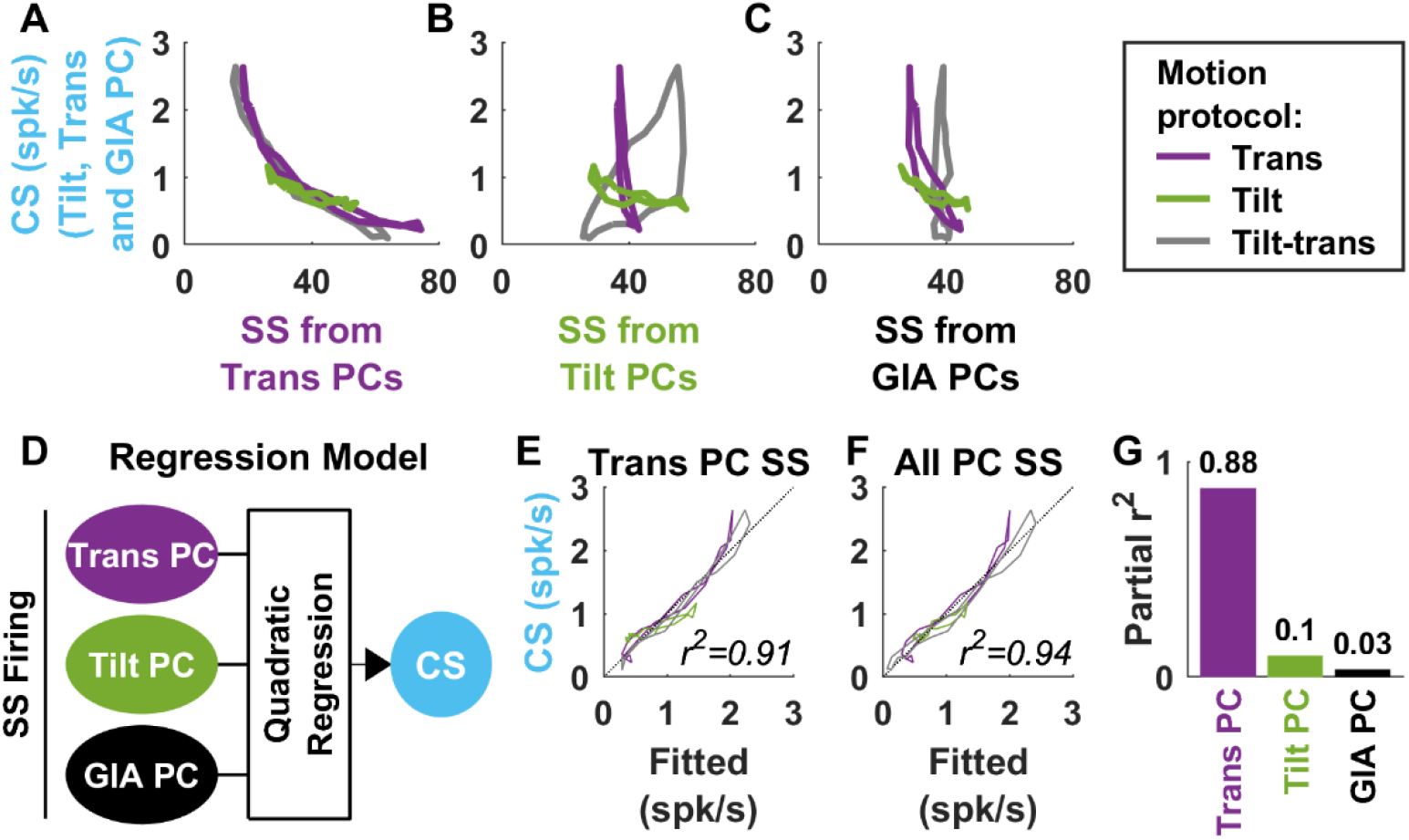
CS firing can be predicted based on SS from translation-selective cells. **A-C:** CS response (from **Fig. 5B-D**; averaged across tilt-, translation- and GIA-selective PCs) versus SS response of translation-selective (A), tilt-selective (B) and GIA-selective PCs (from **Fig. 5B**). There is a clear inverse relationship between CS and the SS from translation-selective cells (A). Importantly, this relationship holds across all motion protocols, including tilt (green). In contrast, there is no consistent relation between the average CS and the average SS responses of tilt- and GIA-selective cells. Therefore, CS firing is predicted based on the SS firing of translation-selective cells only. **D:** To test this, and investigate the SS of tilt-of GIA-selective cells can make any significant contribution to predicting CS firing, we performed a multiple regression analysis where SSs are the predictors and CS the dependent variable. We used a quadratic regression to account for the curvature of the curves in A. **E,F:** Relation between fitted (abscissae) and measured (ordinate) CS firing when the regression uses the SSs of translation-selective cells (E) or of all cells (F) as a predictor. The high r^2^ score in (E) indicates that SS from translation-selective cells explain CS firing accurately, and increases only marginally in (F), indicating and SS from other cell types contributes little. **G:** Partial correlation analysis: based on the same rationale as panels E-F, each variable’s partial r^2^ reflects how much adding this variable to the others increase the regression’s overall r^2^. The partial r^2^ of SSs from translation-selective PC is high and significant (p<10^-3^, shuffling test), whereas the partial r^2^ of SSs from other cell types is not significantly higher than expected by chance (p=0.064 and p=0.41). Thus, from a statistical point of view, the SSs of tilt- and GIA-selective cells don’t contribute significantly to predicting CS firing.

PCs project indirectly to the IO (**Fig. 1A**), and therefore their SS output may take part in controlling IO activity. We performed a regression analysis to evaluate the extent to which CS activity correlates with SS from different groups of PCs. In most cell groups (translation-, tilt- and GIA-selective PCs), CSs occur predominantly during translation, and correlate with the SS firing of translation-selective cells. As CSs and translation-selective cells also modulate to a limited extent during tilt (**Fig. 5B**), it is possible that translation-selective cells alone can predict IO modulation. Indeed, a multiple regression analysis between SS and CS firing (**Fig. 5D**) using a quadratic model to account for the curvature of the curves in **Fig. 5A** demonstrated that the SS activity of translation-selective PCs predicts CS modulation during both translation and tilt, and that adding the SS activity of tilt-selective cells as predictors didn’t improve the fitting significantly (**Fig. 5E-G**).

Thus, in summary, the data in **Fig. 5** offers a synthetic overview of a cell population which putatively constitute a unit of computation in the NU. We will explore the possible architecture of such a circuit further using modelling. But first we will further emphasize the experimental findings by examining CS responses during 3D motion protocols used in (Laurens et al., 2013a, 2013b).

### Three-dimensional responses and motion illusions

We first analysed CS responses of PCs during Tilt While Rotating (TWR; **Fig. 6**; see (Laurens et al., 2013a) for details). TWR consists of alternating tilt (i.e. forward/backward as illustrated in **Fig. 6A**, orange arrows) superimposed on a constant-velocity yaw rotation (**Fig. 6A**, blue arrow). Due to the physical properties of the semi-circular canals, these movements induce an additional rotation signal in a direction orthogonal to the actual tilt (**Fig. 6A**, green arrows). Thus, from a sensory perspective, TWR induces a sideward rotation signal, but without the corresponding activation of the otoliths since the head is in fact immobile in this direction. The only coherent interpretation is that the head undergoes a motion similar to a tilt-translation stimulus (**Fig. 2, 5**) where it tilts and translates simultaneously. Accordingly, TWR induces an illusion of translation (sideward, **Fig. 6A**, violet). In (Laurens et al., 2013a), we demonstrated that the SS firing of translation-selective PCs increases or decreases when TWR induces illusory translation along or opposite to their PD.

**Figure 6:**
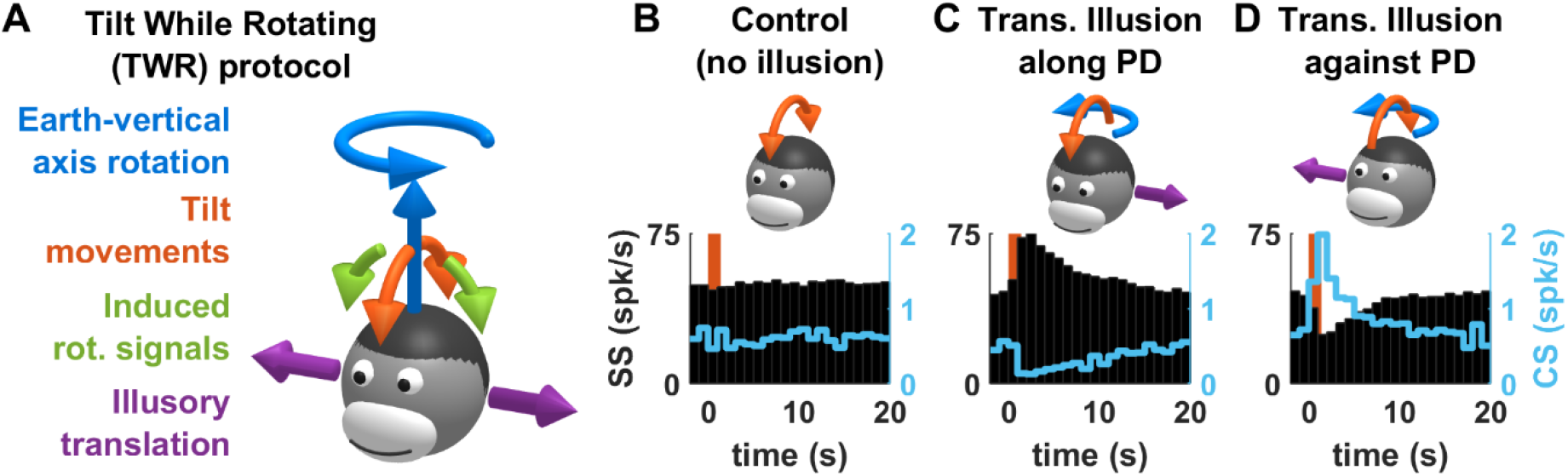
SS and CS responses of translation-selective cells induced by Tilt While Rotating. **A:** Rationale of the protocol (see text for details). **B-D:** SS (black) and CS (cyan) responses during control tilt (B) or TWR inducing illusory translation along (C) or opposite (D) to the SS PD.

We analysed the CS firing of a subset of 9 translation-selective cells in which CSs could be reliably identified. As reported previously (Laurens et al., 2013a), there is no SS modulation when the tilt movement occurred in the absence of yaw rotation (**Fig. 6B**, black). In contrast, there is a strong SS firing increase/decrease when the simultaneous TWR stimulation induced illusory translation along/opposite the cells’ PD (**Fig. 6C,D**, black). The modulation of CSs followed a reciprocal pattern (**Fig. 6C,D**, cyan), such that CSs increased when TWR induced an illusory translation opposite to the cell’s PD. Thus, similar to SS, CS responses are identical during real or illusory motion of the head.

We also analysed CS responses during another illusion-generating motion stimulus, Off-Vertical Axis Rotation (OVAR; **Fig. 7**; see (Laurens et al., 2013b) for details). OVAR consists in tilting the head’s vertical axis (**Fig. 7A**, blue) away from vertical and then rotating at a constant speed about that axis. OVAR causes the head to tilt dynamically (**Fig. 7B**), and can be used to demonstrate that Purkinje cells track tilt accurately when the head rotates about its vertical axis (Laurens et al., 2013b). Furthermore, due to the canal’s high-pass filter properties, angular velocity signals fade out in ~20 s during OVAR. In this situation, accurate tilt perception is gradually replaced by a translation illusion (Vingerhoets et al., 2006, 2007) (**Fig. 7B**, bottom). In (Laurens et al., 2013b), we demonstrated that the SS firing of tilt- and translation-selective cells match the time course of this motion sensation. Indeed, the modulation of tilt-selective cells decreases (**Fig. 7C**) while the modulation of translation-selective cells increases (**Fig. 7D**) as illusory translation builds up.

**Figure 7:**
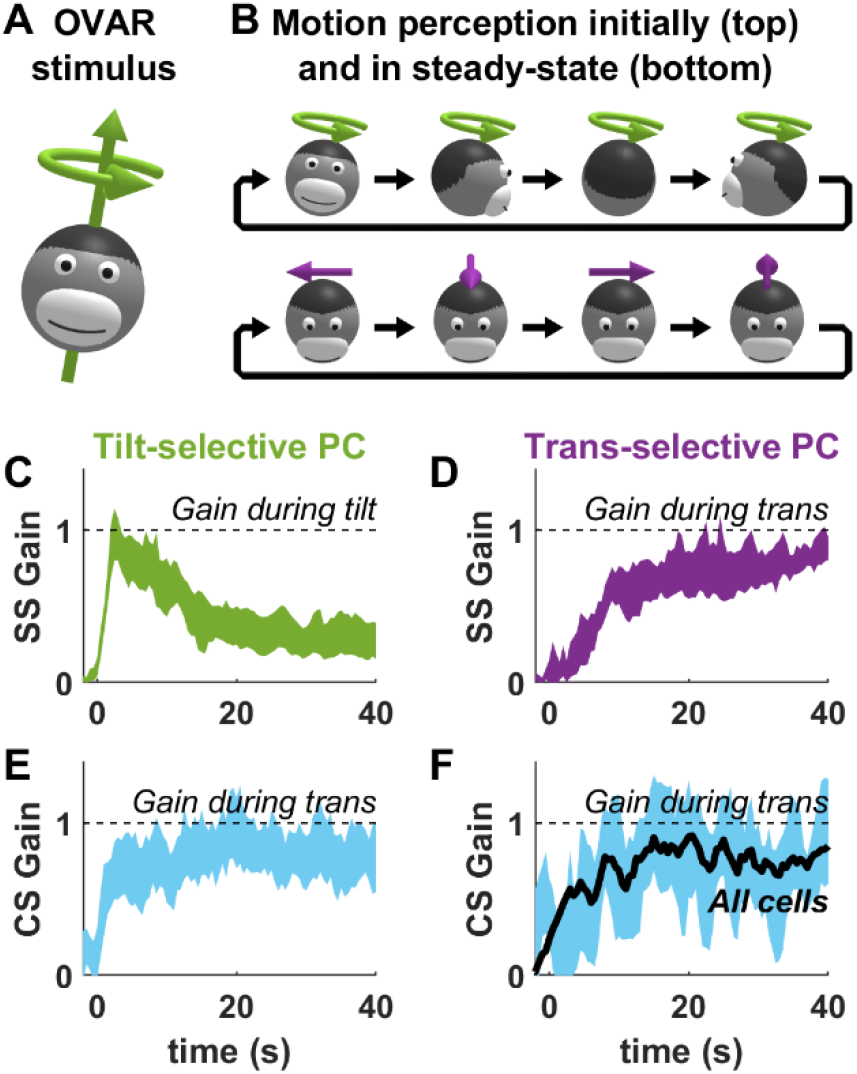
CS responses during OVAR. **A,B:** Rationale of the OVAR protocol. The head rotates at a constant velocity around a tilted axis. Initially (B, top), the motion is perceived veridically. However, rotation signals from the semicircular canals fade out in about 20 s, resulting in a steady-state of an illusion of translation (B, bottom). **C,D:** SS modulation gain of tilt-(C) and translation-(D) selective cells, expressed relative to a tilt stimulus (as in panel B, top) for tilt-selective cells or a translation stimulus (as in panel B, bottom) for translation-selective cells. The bands represent the 95% confidence interval around the mean value. **E,F:** Cyan: CS modulation gain of tilt-(E) and translation-(F) selective cells, expressed relative to a translation stimulus (as in panel B, bottom). The bands represent the 95% confidence interval around the mean value. The black line in F represent the average CS modulation gain of all tilt- and translation-selective cells, pooled.

If indeed CSs occur during real or illusory translation, we expect that both tilt- and translation-selective cells would fire CSs during the late stages, but not the beginning, of OVAR. Further, such a finding would strongly support the hypothesis that CS firing reflects the output of a 3D model of head motion. To test these predictions, we computed the CS modulation gain and phase in both tilt-selective cells (n=14) and translation-selective cells (n=6). We found indeed that CS modulation was low at the beginning of OVAR and increased until it reached a steady-state in both cell types (**Fig. 7, E,F**). When pooled across all cells (**Fig. 7F**, black), the CS response was similar to the SS response of translation-selective cells (**Fig. 7D**).

Collectively, the results of **Fig. 3–7** demonstrate that CS firing in the NU resembles the SS activity of translation-selective cells during 3D motion and illusory motion. This confirms that CS firing is controlled by neurons that implement 3D internal model computations, as outlined in **Fig. 1D**, and supports the possibility that CS are driven by projections from translation-selective cells onto the IO. Considering that these cells encode a feedback signal when sensory prediction errors occur, and that CS firing is involved in cerebellar learning (Ito, 2006; Kimpo et al., 2014; Lisberger, 1988; Nguyen-Vu et al., 2013), the CS recorded in the NU may participate to a learning mechanism triggered by sensory prediction errors.

### Neuronal network model

The finding that both tilt-selective and translation-selective cells receive similar IO inputs support the hypothesis that they may use the same teaching signal to learn two fundamentally different operations. To test whether this hypothesis is computationally feasible, we designed a neural network model (**Fig. 8A,B**) that reflects the computations outlined in **Fig. 1D**, as described below.

**Figure 8:**
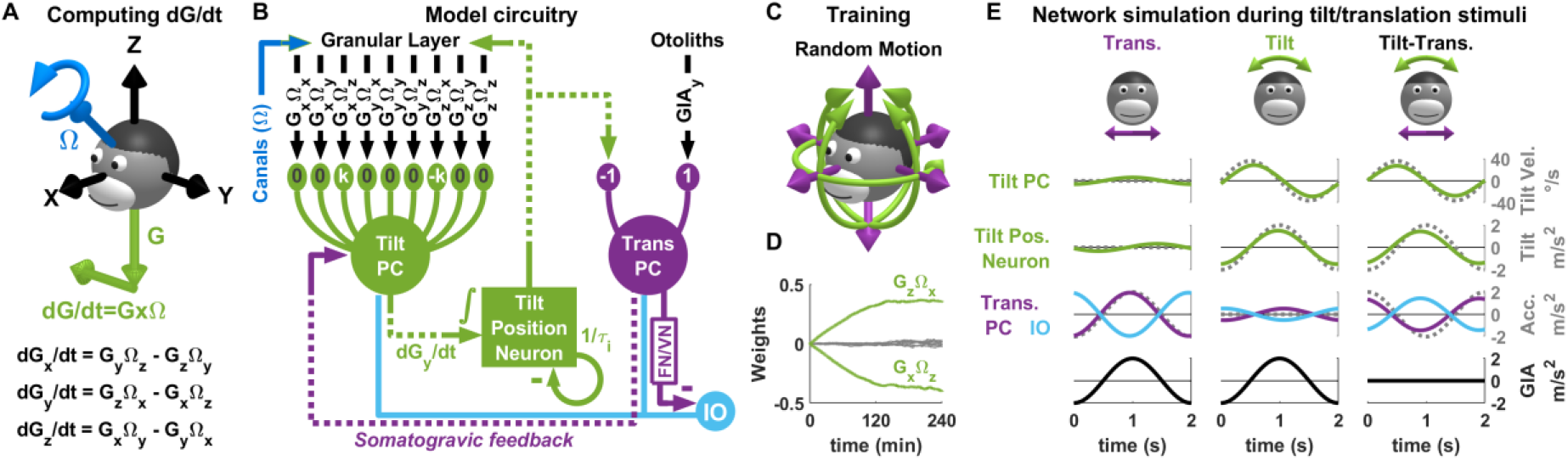
Neuronal network model of tilt/translation disambiguation. **A:** Mathematical formulation of the transformation performed by tilt-selective cells. Converting egocentric rotation signals (Ω blue) into tilt-velocity (dG/dt, green) requires a vectorial cross-product, whose formula is shown at the bottom of the panel. **B:** Structure of one olivo-cerebellar loop in the simulated neuron network. This loop receives otolith inputs (top right) encoding the lateral component of the GIA, i.e. GIAY. During learning, the PD of all other cells align with this axis. In the full network, we simulate loops receiving otolith inputs along all cardinal axes. These loops operate independently, with one exception: the granular layer upstream of tilt PCs receives output from tilt position neurons from all loops, which provide the gravity signal G along all dimensions. **C:** The network is trained using random rotations and translations in 3D. **D:** Evolution of the synaptic weights of a tilt PC encoding dGY/dt. Bands represent mean ± sd over 15 simulations. Green: synaptic weights of the components required to compute dGY/dt; grey: synaptic weights of other components. **F:** Simulated response of all neurons in the network during tilt, translation and tilt-translation.

Tilt-selective PCs (**Fig. 8B**, ‘Tilt PC’) compute tilt velocity, i.e. the derivative of gravity dG/dt. Mathematically, tilt velocity can be expressed as the vectorial cross-product GxΩ, which can be decomposed into combinations of products (**Fig. 8A**): for instance, lateral tilt velocity, dG_Y_/dt, is computed as G_Z_Ω_X_ – G_X_Ω_Z_. We propose that granule cells encode all 9 possible products (G_X_Ω_X_, G_X_Ω_Y_, etc; **Fig. 8B**), and that tilt-selective PCs learn to combine these products. For instance, to encode dG_Y_/dt (with a gain factor k), a tilt-selective cell would associate a weight of k to G_Z_Ω_X_, -k to G_X_Ω_Z_, and 0 to all other products. In this respect, our model follows the Marr-Albus hypothesis, in which granule cells act as basis functions (Albus, 1971; Marr, 1969). In (Laurens et al., 2013b; Laurens & Angelaki, 2020), we noted that, even though tilt-selective PCs primarily encode tilt velocity, their firing is partially shifted toward tilt position. To reproduce this property, we modelled these cells as leaky integrators with a short time constant of 50ms.

The output of tilt-selective PCs is integrated temporally to yield an estimate of tilt position (i.e. G) by “tilt position neurons” (whose nature is currently unknown) (**Fig. 8B**). Since it is questionable whether neurons can perform a perfect integration, we model these neurons as leaky integrators with a time constant of 1.2s. Tilt position neurons project to translation-selective PCs, as well as to the granular layer in which they provide the gravity signal required to compute GxΩ.

Translation-selective PCs (**Fig. 8B**) combine the tilt estimate and the raw otolith signals to compute net translation. The polarity of this connection can be deduced as follows: based on **Fig. 5**, we know that a translation cell with a given PD (e.g. leftward acceleration) receives the same IO input as a tilt-selective cell with an equivalent PD (e.g. rightward tilt velocity, see **Fig. 1B**). Therefore, the pathway between tilt-selective and translation-selective PCs should be altogether *inhibitory*. Note, however, that the actual pathway between tilt-selective and translation-selective cells involves an unknown number of synapses. Here we describe it as an entirely excitatory pathway that terminates with a final inhibitory synapse to the translation-selective PC. In practice, the polarity of the individual connections may be changed without loss of generality.

Translation-selective cells also project to tilt-selective cells (**Fig. 8B**; see also **Fig. 1D**), as shown in (Laurens et al., 2013b) to implement a well-known mechanism (Graybiel, 1952) called somatogravic feedback (Laurens & Angelaki, 2011, 2017), which prevents the tilt position neurons from accumulating errors as they integrate noisy inputs over time and compensates for the leaky dynamics of the tilt position neurons.

### Learning rule

Next, we assume that IO signals drive synaptic plasticity according to the following rule (Dean et al., 2002, 2010; Dean & Porrill, 2014):

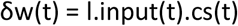

Where input(t) and cs(t) are the synaptic inputs for a given synapse and the IO inputs, respectively, δw(t) is the change of synaptic weight, and l a learning factor. This rule implements a mechanism called ‘decorrelation learning’ (Dean et al., 2002, 2010; Dean & Porrill, 2014) through which the circuit outlined in **Fig. 8B** learns to cancel its otolith input (GIA) based on its canal input (Ω), which amounts to computing a tilt signal. Note that, in this model, this mechanism is more elaborate than in previous work (Dean et al., 2002, 2010; Dean & Porrill, 2014) since it involves a non-linear operation (the cross-product GxΩ) as well as a temporal integration from tilt velocity to position.

A priori, synaptic plasticity could occur at all synapses in the network. However, in the simulated network, any change in the synaptic weight between tilt-selective PCs and tilt position neurons, or between tilt position neurons and translation-selective PCs, can be replaced by an overall gain change of the active synapses of tilt-selective PC. Therefore, for simplicity, we only consider plasticity at the level of tilt-selective PCs.

### Simulation

In order to create a full 3D model, we simulated 3 parallel loops that process GIA_X_, GIA_Y_ and GIA_Z_. The tilt position neurons in these 3 loops provide the components G_X_, G_Y_, G_Z_ required to compute GxΩ. We trained the model during simulated 3D motion (**Fig. 8C,D**), and then simulated all cell types during roll tilt and lateral translation (**Fig. 8B**). The synaptic weights of the tilt-selective PC were initialized randomly (following a Gaussian distribution with a standard deviation of 0.2) priori to training. During training, the weights corresponding to G_Z_Ω_X_ and G_X_Ω_Z_ evolved in opposite directions and stabilized to opposite values (**Fig. 8D**, green), while all other weights decreased to 0 (**Fig. 8D**, grey). This indicates that the synapses of the simulated tilt-selective cells implement the cross-product necessary to compute dG_Y_/dt.

The simulated neuronal responses during tilt/translation paradigms reproduced the prominent properties of tilt- and translation-selective PCs. Tilt-selective PCs responded during tilt (**Fig. 8E**, middle) with a gain of 0.78 relative to tilt velocity and were primarily in phase with tilt velocity, but shifted by 11° towards tilt position. During translation (**Fig. 8E**, left), their response gain was largely reduced (4.6 times less than during tilt). Tilt position neurons also responded during tilt specifically, with a gain of 0.76 and a slight phase lead of 5° relative to tilt position (**Fig. 8E**, middle). Their response to translation (**Fig. 8E**, left) was also much lower (by a factor of 4.8 compared to tilt). In contrast, translation-selective cells responded during translation (**Fig. 8E**, left) with a gain of 0.97 and phase lead of 9°, and their response during tilt was reduced by a factor of 3.8 (**Fig. 8E**, middle). The simulated IO response was the inverse of that of translation-selective PCs. As expected, all cells responded during tilt-translation, and maintained their phase relative to tilt velocity and position (tilt PC and tilt position neurons) or translation (translation PC and IO neurons).

Thus, a simple CS-driven learning rule, based on the principle of decorrelation learning, where only translation-selective cells project to the IO, is sufficient to train a neuronal network to integrate rotation signals in 3D so as to predict and cancel tilt-driven activation of the otoliths. These computational models support the experimental finding that a single olivo-cerebellar loop with a shared error signal underlies the diverse vestibular modulation encountered in the vermal vestibulocerebelum.

## Discussion

Olivo-cerebellar loops form a unit of cerebellar computation (Apps et al., 2018; Apps & Garwicz, 2005; Chaumont et al., 2013; De Zeeuw et al., 2011; Ekerot et al., 1991; Garwicz et al., 1998; Herzfeld et al., 2015; Ozden et al., 2009; Shadmehr, 2020; Sugihara & Quy, 2007). Here we show that two functionally distinct types of PC may implement two computational steps within a single olivo-cerebellar loop.

In previous studies (Angelaki et al., 2004; Laurens et al., 2013b; Laurens & Angelaki, 2020; Yakusheva et al., 2007, 2008, 2010), we have identified two distinct groups of PCs defined by their SS properties: Tilt-selective cells encode allocentric velocity relative to gravity which, when integrated, can predict the gravitational force acting on the inner ear’s inertial sensors - the otoliths. Translation-selective cells encode otolith prediction error. Yet, the CS properties of both cell types are identical and proportional to the SS firing of translation-selective cells, i.e. to the otolith prediction error. This finding suggests that translation-selective PCs may control the activity of IO cells that innervate them (Chaumont et al., 2013) through their downstream projections to the fastigial or vestibular nuclei. Thus, the output of translation-selective PCs may serve as a dual function - driving behavioural responses and generating a teaching signal to maintain optimal control through the IO loop.

The similarity in CS response properties suggests that both types of PCs belong to a single olivo-cerebellar loop. Thus, tilt- and translation-selective cells may form a computational unit that uses its own output as a teaching signal. In this respect, they may implement the decorrelation learning rule proposed by (Dean et al., 2002, 2010; Dean & Porrill, 2014) to explain how efference copies are used to filter out self-generated actions from a sensory signal. The computations performed by tilt- and translation-selective cells are, however, more intricate than the reafference suppression function because they involve a 3D non-linear spatial transformation combined with temporal integration. Indeed, our model simulations confirm that such computations can be learned using a decorrelation learning rule.

### Tilt- and translation-selective cells form a computational unit

Previous studies (Angelaki et al., 2004; Laurens et al., 2013b; Laurens & Angelaki, 2020) have shown that tilt- and translation-selective cells encode the two interconnected computational steps outlined in **Fig. 1C**. Yet, their functional link had remained tentative without any established neural pathway between tilt-selective and translation-selective PCs. Alternatively, it could be that these properties arise independently in these PC types, perhaps through computations that occur elsewhere, e.g., in the granular layer or the vestibular nuclei. The current finding that both cell types receive identical IO inputs supports the notion that they are functionally linked within the same olivo-cerebellar network. Our findings also provide answers to the following question: if a neuronal pathway links tilt-selective and translation-selective PCs, then, considering that this pathway is likely polysynaptic, is it overall excitatory or inhibitory? For instance, a tilt-selective cell whose SS firing encodes leftward tilt (after temporal integration) may either inhibit a translation-selective cell that encodes rightward acceleration (**Fig. 1B-D**) or activate a translation-selective cell that encodes leftward acceleration. Our finding that cells that prefer e.g. leftward tilt and rightward acceleration would receive identical IO inputs (**Fig. 5**) supports the former possibility, and suggests that the postulated anatomical link is overall inhibitory.

### Internal model computations for self-motion perception and feedback signals

The concept of internal model is a classical approach for apprehending how the brain processes multisensory self-motion information, proposed as early as the late 70s (Oman, 1982; Ormsby & Young, 1977). Several quantitative models were subsequently developed in the following decades (Borah et al., 1988; Bos & Bles, 2002; Glasauer & Merfeld, 1997; Karmali & Merfeld, 2012; Laurens & Angelaki, 2011, 2017; Laurens & Droulez, 2007; Merfeld, 1995; Zupan et al., 2002), whose findings have been extensively validated by behavioural (Angelaki et al., 1999; Dakin et al., 2020; Khosravi-Hashemi et al., 2019; Laurens et al., 2010, 2011; Merfeld, 1995; Merfeld et al., 1999) and neurophysiological studies (Angelaki et al., 2004; Cullen, 2012; Cullen & Brooks, 2015; Cullen & Roy, 2004; Hernández et al., 2020; Laurens et al., 2013a, 2013a; Laurens & Angelaki, 2020; Shaikh et al., 2005; Stay et al., 2019; Yakusheva et al., 2007, 2008, 2013).

Whereas early studies have focused on passive motion (Angelaki et al., 2004; Laurens et al., 2013a, 2013a; Laurens & Angelaki, 2020; Shaikh et al., 2005; Yakusheva et al., 2007, 2008, 2013), we have proposed a more general framework (Laurens & Angelaki, 2017) in which cerebellar PCs implement a forward model of the otolith organs, and in which translation-selective cells encode the resulting sensory prediction error.

This theoretical hypothesis has already been supported by multiple experimental findings. First, translation-selective cells respond to passive translation. Second, their firing is reduced when other sources of information can be used to predict otolith activity; e.g., efference copy signals during active translations; indeed, the firing of vestibular and fastigial nuclei translation-selective cells is markedly reduced (Carriot et al., 2013; Mackrous et al., 2019) - a finding which presumably generalises to translation-selective PCs in the NU. Further, responses of translation-selective cells is also diminished during tilt, during which rotation signals can be used to track head tilt relative to gravity and predict the gravitational activation of the otoliths (Glasauer & Merfeld, 1997; Laurens & Angelaki, 2017; Merfeld, 1995). Finally, this framework implies that an otolith prediction error, and a corresponding activation of translation-selective cells, should occur whenever rotation signals do not match head motion relative to vertical. We have verified this hypothesis in (Laurens et al., 2013a, 2013b), and shown that the firing of translation-selective neurons reflect these illusory signals (Laurens & Angelaki, 2011).

### Neuronal and behavioural outputs of the NU

Translation-selective cells have been identified in multiple brain areas: the fastigial and vestibular nuclei (Angelaki et al., 2004; Hernández et al., 2020; Laurens et al., 2013a, 2013a; Laurens & Angelaki, 2020; Shaikh et al., 2005; Stay et al., 2019; Yakusheva et al., 2007, 2008, 2013) and the vestibular thalamus (Dale & Cullen, 2017).

In contrast, tilt-selective cells have, to date, only been formally identified in the NU. This may be because identifying cells that encode rotation velocity relative to vertical requires distinguishing them from semicircular canal-driven cells that encode rotation velocity in egocentric coordinates. Tilt-selective cells and canal-driven cells have similar responses during simple rotations about earth-vertical or earth-horizontal axes, as in e.g. **Fig. 2,5**. Therefore, formally identifying tilt-selective cells requires testing their responses during multiple 3D rotation protocols, e.g., as in **Fig. 7**, which has only been done in the NU so far (Laurens et al., 2013b). For example, some rotation-selective neurons in the vestibular and fastigial nuclei (Buettner et al., 1978; Büttner et al., 2003; Siebold et al., 1997; Waespe & Henn, 1979) have been presented as tilt-selective (Mackrous et al., 2019), but it is currently unknown whether these cells encode tilt, as opposed to egocentric rotation. Note that tilt signals have been identified in the navigation system (Angelaki et al., 2020), but these cells were not tested during tilt/translation discrimination protocols, thus it is unknown whether they convey a net tilt signal. Although tilt perception is driven by a 3D internal model in humans (Clark & Graybiel, 1966; Merfeld et al., 2001; Niehof et al., 2019a, 2019b; Vingerhoets et al., 2007), whether tilt-selective cells exist outside the NU remains unknown.

Beyond self-motion perception, the NU innervates regions of the fastigial nucleus that are involved in attention, vigilance and hippocampal function (Fujita et al., 2020), suggesting a possible consequence of otolith prediction errors for a variety of brain functions.

### Anatomical substrate of the olivo-cerebellar loop with the NU

Olivo-cerebellar loops in the NU have been studied in rabbits (Barmack & Shojaku, 1995; Fushiki & Barmack, 1997), mice (Yakhnitsa & Barmack, 2006) and cats (Kitama et al., 2014) using exclusively rotation (but not translation) stimuli. Although it is impossible to test the current hypotheses in the absence of translation stimuli, findings from these studies are consistent with the present results. First, these studies reported SS and CS modulation during tilt, but not during rotations in an earthhorizontal plane. Note that even translation-selective cells show a substantial modulation during tilt (**Fig. 3A**, **Fig. 5**). It is thus possible that neurons recorded in previous studies reflected a mixture of tilt-selective and translation-selective cells. In fact, one study (Kitama et al., 2014) noted that NU cells could respond in phase with either tilt position or velocity. Considering that tilt-selective cells encode velocity (Laurens & Angelaki, 2020), ‘velocity’ cells likely correspond to tilt-selective cells, whereas ‘position’ cells likely correspond to translation-selective cells. This interpretation is corroborated by the SS modulation gain during tilt at 0.5 Hz: 133 and 64 spk/s/G respectively for ‘velocity’ and ‘position’ cells respectively in (Kitama et al., 2014), that match our recordings (**Fig. 3A**). Furthermore, previous studies also found that CSs occur in antiphase with SSs, in agreement with our observations (**Fig. 5**). Altogether, these similarities indicate that the NU PCs reported to be modulated by tilt in (Barmack & Shojaku, 1995; Fushiki & Barmack, 1997; Kitama et al., 2014; Yakhnitsa & Barmack, 2006) likely correspond to both tilt- and translation-selective cells.

The medial portion of the NU receives projections from two regions of the IO. The first, which is composed of the dorsal cap and ventrolateral outgrowth, carries visual optokinetic signals (Barmack & Hess, 1980; Leonard et al., 1988) to a small medial portion of the nodulus. Since our experiments were performed in darkness, this region is unlikely to account for the CS responses studied here. The second IO region is the beta nucleus (Barmack, Fagerson, Fredette, et al., 1993; Voogd et al., 1996), which receives projections from the medial and descending vestibular nuclei (Balaban & Beryozkin, 1994; Barmack, Fagerson, & Errico, 1993; Gerrits et al., 1985; Saint-Cyr & Courville, 1979; Turecek & Regehr, 2020) and the parasolitary nucleus (Barmack, 2006; Barmack & Yakhnitsa, 2000). In turn, the medial and descending vestibular nuclei receive projection from the NU (Bernard, 1987; Epema et al., 1985; Shojaku et al., 1987; Wylie et al., 1994), as well as the parasolitary nucleus (Barmack, unpublished observations reported in (Barmack & Yakhnitsa, 2000), and R. Sillitoe, personal communication). Thus, the anatomical substrate of the olivo-cerebellar loop involving tilt and translation-selective cells may include a projection of the NU to the beta nucleus through the medial and descending vestibular and parasolitary nuclei. In agreement with this hypothesis, translation-selective cells exist in the vestibular nuclei (Angelaki et al., 2004; Zhou et al., 2006); the firing of neurons in the parasolitary and beta nuclei is similar to the CS responses in the NU (Barmack, Fagerson, Fredette, et al., 1993; Barmack & Yakhnitsa, 2000), and Fos expression studies indicate that the beta nucleus is activated by linear accelerations (Li et al., 2013).

### Conclusion

From an experimenter’s point of view, linking neural circuits and theoretical predictions may appear an arduous if not vain undertaking, since abstract concepts such as internal models and Kalman filtering may seem too far remote from physiological reality, if not plainly “too nice”. Indeed, we too are amazed that the vestibulo-cerebellar circuit should consistently reflect such theorized computations. And yet, these findings should not come as a surprise, since behavioural studies have consistently shown that the brain implements the building blocks of internal models, which are nicely mathematically tractable in a well-defined computational challenge such as tilt/translation discrimination. Thus, when theoretical concepts have passed the test of decades of scrutiny, we should expect to find their embodiment in neuronal circuits.

## Acknowledgements

The work was supported by NIH grant DC004260.

## Methods

### Animals

Three male rhesus Macaques, aged 3, 4 and 9 years, were used in the study. The animals were pair-housed in a vivarium under normal day/night cycle illumination. Experimental procedures were in accordance with US National Institutes of Health guidelines and approved by the Animal Studies Committee at Washington University in St Louis (approval n°20100230).

### Experimental procedures and neuronal recordings

Experimental procedures were described in detail in (Laurens et al., 2013a, 2013b). In summary, animals were seated in primate chairs that were installed on a 3-axes rotator mounted on a linear sled (Acutronics Inc, Pittsburg, PA). We recorded neurons extracellularly using expoxy-coated tungsten electrodes (5 or 20 MΩ impedance; FHC, Bowdoinham, ME). Recording locations were determined stereotaxically and relative to the abducens nucleus. Raw spiking data was sorted offline using custom Matlab scripts, based on spike amplitude and principal components analysis. In this study, we included only neurons where CS firing could be isolated consistently across trials, and where CS were followed by a pause in SS firing for at least 10 ms.

### Experimental protocols

Sinusoidal tilt and translation stimuli (**Fig. 2A**) consisted of translation (peak acceleration = 0.2 g, with g = 9.81 m/s^2^) or tilt (peak tilt = 11.5°) oscillations at 0.5 Hz, or combinations of these stimuli (out of phase: tilt-translation or in phase: tilt+translation). Stimuli could be delivered along the head’s naso-occipital axis (forward/backward translation and pitch tilt), lateral axis (left/right translation and roll tilt) or along intermediate axes. We recorded the responses of each cell using stimuli along at least two head axes.

Tilt while rotating (TWR) (Laurens et al., 2013a) consists of rotating the setup about a fixed earth-vertical axis at a constant velocity of 45°/s. During this rotation, animals were tilted back and forth ±10° along one plane (i.e. pitch, roll or intermediate) about the vertical axis. Tilt movements were brief movements (peak velocity 20°/s, acceleration 50°/s^2^, duration 1.4s) that were separated by 30s of fixed tilt.

Off-vertical axis rotation (OVAR) (Laurens et al., 2013b) consisted in tilting the animal by 10°, and then rotating around the head’s vertical axis at 180°/s (peak acceleration: 90°/s^2^) for 80s. This resulted in the head tilting in a sequence (nose up, left ear down, nose down, right ear down, nose up) which is physically equivalent to out-of-phase oscillations in pitch and roll, with 10° peak tilt, at 0.5 Hz.

### Data analysis

SS and CS firing were analysed using the same methods as in (Laurens et al., 2013a, 2013b). We also refer the reader to (Laurens & Angelaki, 2016) for an in-depth presentation of the analysis of sinusoidal tilt and translation stimuli. In this section, we present some analyses that were specifically developed or modified in the present study.

#### Modulation amplitude

The only difference between the analysis of SS and CS was the way modulation amplitude was computed. To quantify the modulation of SS, we fitted firing histograms with a rectified sinusoid function: FR(t) = max(0;FR_0_+A.cos(π.ω.t+ϕ)) where ω is the stimulus frequency in Hz, A and ϕ the response amplitude and phase and FR_0_ the cell’s baseline firing. To quantify the modulation of CS, we performed a simple Fourier transform, which is equivalent to fitting firing histograms with a sinusoid FR(t) = FR_0_+A.cos(π.ω.t+ϕ), without rectification. We chose this approach because using a rectified function yields more accurate results for cells where the firing becomes ‘less than 0’ in the trough of the firing histograms, but is unreliable when cells discharge a low number of spikes, which is the case with CS.

#### Response PD and phase

In **Fig. 4**, we summarize the cells’ firing properties by computing the PD and response phase of SS and CS. For instance, a cell may respond to leftward acceleration with a phase lead of 10°. However, it is equivalent to this cell responding to rightward acceleration with a phase lead of −170°. In order to express the PD and phase of SS and CS in a coherent manner, we adopt the following procedures. First, we compute the PD and phase of SS. For translation-selective cells, we express the PD such that the response phase during translation is within ±90° (in the example above, the PD would be to the left). For tilt-selective cells, we express it such that the response phase during tilt is always within 54±90°: this is because tilt-selective cells encode a mixture of tilt velocity and position, with a mean response phase of 54° at the population level (Laurens et al., 2013b), and therefore it is logical to express response phase in an interval centred around that value. For other cell types, we proceed as for translation-selective cells. Note that this convention has no impact on our statistical analyses but only serves to make figures clearer (e.g. in **Fig. 4C**).

Next, we compute the PD and phase of CS, independently from the SS response. In a second step, if the PDs of SS and CS are more than 90° apart, we invert both the PD and phase of CS. In absolute terms, these conventions do not change how we measure the SS and CS responses, since reversing both the PD and phase results in an equivalent description of the response. Note that, using these conventions, (1) the absolute difference between the PD of SS and CS is always less than 90° (**Fig. 4A,B**); (2) the phase of SS is expressed as a circular variable with a periodicity of 180°; whereas (3) the phase of CS is a circular variable with a periodicity of 360° (**Fig. 4C,D**).

#### Regression analysis

We tested whether CS firing can be related to the SS output of PC populations by performing a multiple regression analysis, where the dependent variable was the average CS firing in the NU (*CS_NU_*) and the predictors were the SS firing of translation-, tilt- and GIA-selective cells (*SS_trans_*, *SS_tilt_* and *SS_GIA_*). We constructed a vector *SS_trans_* that contains the SS firing histograms of translation-selective cells during translation, tilt and tilt-translation (i.e. the three black histograms in Fig. 5A, concatenated one after another). We constructed the vectors *SS_tilt_* and *SS_GIA_* similarly. Finally, we constructed a similar vector *CS_NU_* by averaging the CS firing of all translation-, tilt- and GIA-selective cells. Next, we build a quadratic regression model:

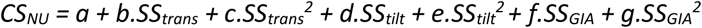

We evaluated the goodness of fit of the regression by computing the squared coefficient of correlation R^2^ = 1-SSR/SS_tot_ where SSR is the sum of squared residuals and SS_tot_ the variance of *CS_NU_*. To measure the contribution of each SS response type, we computed partial R^2^, e.g. for translation-selective cells, we computed pR^2^_trans_ = 1-SSR/SSR_trans_, where SSR_trans_ is the sum of squared residuals obtained when translation-selective cells are excluded from the regression. A large/small pR^2^ indicates that including a given response type has a large/small impact on the regression’s goodness of fit, implying that SS from the corresponding population of PC contribute to a large/small extent to controlling CS firing. We used a shuffling approach to estimate the confidence intervals of pR^2^: we computed 10000 shuffled values of pR^2^_trans_, for each of which the vector *SS_trans_* was shuffled, and defined the confidence interval (at α = 1%) as the 99-percentile of the distribution of shuffled values. We performed the same computation for pR^2^_tilt_ and pR^2^_GIA_. We found that the 99-percentile is equal to 0.16 in all cases, i.e. pR^2^ values lower than 0.16 were not significantly different from 0.

## Notes

### Competing Interest Statement

The authors have declared no competing interest.

